# Storage polymer polyhydroxybutyrate promotes recovery and limits phenotypic heterogeneity in regrowth from carbon starvation in *Methylobacterium extorquens*

**DOI:** 10.64898/2025.12.12.690574

**Authors:** Keaton J. Adams, Christopher J. Marx, Alexander B. Alleman

## Abstract

Bacteria in the wild often encounter long periods of starvation. Evolutionary adaptation to cycles of nutrient deficiency has led to metabolic and physiological strategies that prevent cell death and promote rapid recovery from starvation. A cellular factor shaping outcomes during carbon starvation is polyhydroxybutyrate (PHB), a widespread carbon polymer that helps maintain membrane integrity and promotes regrowth upon reintroduction to nutrients. The response to starvation is often highly heterogeneous across isogenic populations and confounded by environmental variables that mask underlying physiological mechanisms. Prior work has shown that hyper- or hypo-osmolarity greatly increases cell death, but little is known about how PHB and osmolarity jointly influence both population viability and the emergence of heterogeneity in periods of nutrient limitation. To investigate this, we subjected populations of *Methylobacterium extorquens* to carbon starvation across a range of osmolarities and analyzed the phenotypes of individuals regrowing from the combined stress. By resolving the distributions of key traits such as lag time and growth rate, we found that populations partition into subpopulations that appear either significantly or minimally impaired by the stresses presented. Osmolarity affected total survival, and surprisingly, populations maintained viability in ultra-pure water. PHB metabolism was beneficial for population viability across all conditions, and unexpectedly reduced the variance of populations in lag time and growth rate. These findings show that PHB aids survival and regrowth from starvation, and suggest that the restructuring of phenotypes in long-term stationary phase depends on PHB even though total osmotic sensitivity does not.

## Introduction

Nutrient influx for microbes in the environment is typically sparse and intermittent. These cycles of feast and famine generate ecological dynamics distinct from those traditionally studied in nutrient-rich media (1, 2). For bacteria less equipped to handle starvation, this can represent formidable stress. The gut bacterium *Escherichia coli* starved in minimal media lacking a carbon source reached 1% of its original viability in just seven days (3). In some taxa, adaptation to famine has uncovered particularly clever strategies, like spore formation, dormancy, or extremely slow growth (4–6). However, reemergence from dormant or spore states takes significant time, and reliance on these strategies for survival in starvation is an energetically costly commitment (7). This presents a key tradeoff between tolerance to famine and the speed of reestablishing growth in feasts, as initiating growth as soon as possible following a nutrient influx is crucial for competition and colonization success.

One primary factor that aids cells in starvation is the accumulation of carbon storage polymers, such as polyhydroxybutyrate (PHB). Cellular PHB synthesis during episodes of carbon excess forms large, opaque, intracellular granules, which bacteria can later depolymerize to produce NAD(P)H and acetoacetyl-CoA to provide basic cell maintenance (8–10). Production of cell PHB content is largely exclusive to carbon excess environments, with nitrogen, metals, or oxygen often limiting growth (11). Typical laboratory carbon to nitrogen (C:N) ratios of around 2:1 lead to average PHB mass per bacterium of 5 percent or less, while higher ratios lead to significantly higher PHB mass (12). Reports of monoculture PHB production record maximum average PHB mass per bacterium of 30-60 percent, with some high-performing strains of *Cupriavidus necator* yielding 80 percent (13, 14). In several α-proteobacterial taxa (Rhizobiaceae, Burkholderiaceae, etc.) commonly found in soil, PHB has been shown to improve viability and regrowth, even enabling a cell to double after removing an exogenous carbon source (15, 16).

External fluctuations in pH, temperature, and osmolarity can also affect survival, frequently overshadowing the effects of nutrient limitation alone (17). Osmotic pressure is such an influential factor that the death rate of *E. coli* in starvation can vary over a 40-fold range simply by changing the salt concentration of the medium. This is due to the fact that *E. coli* spends over 90% of its energy budget in starvation on exporting ions (3). Sensitivity to osmotic environments is varied across taxa: tolerance to hypertonic desiccation is phylogenetically broad, emerging independently in select lineages of Actinomycetota, Firmicutes, and Proteobacteria (18–20), while tolerance to hypoosmotic stress is less reported, and concentrated in Proteobacteria (21–23). The physiological basis of this apparent lineage-specific proficiency is not well characterized, but its phylogenetic asymmetry highlights that osmotic tolerance strategies may rely on fundamentally distinct cellular mechanisms.

Environmental fluctuations also generate variance in growth phenotypes, such as wide lag time distributions in *E. coli* switching resources, or variable expression of virulence regulons in *Salmonella enterica* Typhimurium upon entry to host cells (24–26). Stressors like osmotic pressure exacerbate this variance, as tolerance to stress typically defines the threshold between life and death, and cells will sacrifice performance in other phenotypic traits to maintain survival. This was seen in populations of *Campylobacter jejuni*, where high osmotic pressure led to skewed and multi-modal distributions of lag time in populations that were normally distributed if unstressed (27). For other stressors like formaldehyde, populations of *Methylobacterium extorquens* were observed diversifying into discrete subpopulations with distinct tolerance phenotypes, leading to bimodally distributed colony appearance times when regrown free from stress (28). In non-growth environments as well, heterogeneity provides an adaptive benefit, diversifying bacterial populations into viable but non-culturable (VBNC), persister, and spore states (4, 5, 29, 30). In yeast, glucose starvation generates a small fraction of reversibly dormant cells that are more equipped to handle nutrient limitation (31). Differential or stochastic gene expression directly underlies these instances of phenotypic heterogeneity, and can generate trait distributions that take on a wide variety of shapes, including bimodality, under different regulatory topologies (32–34). These instances of heterogeneity, compounding with environmental pressures, make predicting survival outcomes in starvation challenging.

We sought to investigate the joint effects of carbon starvation and osmotic pressure upon bacteria to better contextualize the long-term survival of bacteria in unfavorable environments. *Methylobacterium*, a genus of methylotrophic α-proteobacteria, provides an ideal model for examining these joint effects. *Methylobacterium* are commonly found in leaf and soil microbiomes of plants, municipal water systems, and even air conditioning evaporator cores (35–37). These habitats are characterized by infrequent nutrient input and frequent osmotic stress, yet *Methylobacterium* maintain robust and diverse communities (38, 39). An underexplored potential mechanism explaining long-term survival in *Methylobacterium* is carbon storage in PHB. However, PHB per cell mass is widely heterogeneous and environment dependent. In carbon-excess conditions, production of PHB varies widely in both number of granules formed and total PHB per cell mass, ranging from 0 to 85 percent in some species (40). It is unclear whether the heterogeneity in PHB production leads to subsequent heterogeneity in survival phenotypes (death rate, lag time, and regrowth rate) for starved populations. Additionally, it is unknown how the beneficial effects of PHB interact with external osmolarity to modulate total viability and the diversification of cell fates during starvation.

To better understand the effects of PHB heterogeneity on survival during starvation, we subjected populations of *M. extorquens* PA1 to carbon starvation across a range of osmotic conditions. We compared wild-type (WT) populations to a mutant unable to synthesize PHB (Δ*phaC*) and analyzed total viability and the distributions of phenotypic traits after starvation stress. The presence of PHB was hypothesized to provide a survival and regrowth benefit, but its potential effect on the variance of populations in lag time and growth rate was less clear. If the beneficial effects of PHB metabolism outweighed the heterogeneity in its production, the presence of PHB would limit variance in lag time and growth rate by promoting equal outcomes from starvation. Alternatively, If PHB mass directly correlated with cell survival, variance in PHB production would directly generate and increase variance in regrowth phenotypes. We determined that PHB enhances survival, critically alters the structure of subpopulations in long-term starvation, and provides a benefit independent of osmolarity.

## Results

### The presence of PHB positively impacts survival in starvation and subsequent regrowth

We first sought to quantify the effect of PHB metabolism upon survival for *M. extorquens* PA1 in a starvation environment, as it has been shown in numerous other model organisms to play a critical role (9, 16, 41). In order to disable PHB synthesis, we generated an unmarked Δ*phaC* strain by allelic exchange (42). Both WT and Δ*phaC* were grown in standard succinate media to stationary phase, washed, and then transferred to media containing no carbon or nitrogen. We assayed viability from colony forming units (CFU) for up to twenty days in starvation conditions. Whereas many bacteria show an immediate loss of viability upon entry into long-term stationary phase (3), *M. extorquens* WT did not significantly decrease in viability over the first 12 days (t-test, p = 0.082 for WT day 0 vs. 12 CFUs) (Fig. 1A). After the first significant drop in viability, loss in CFU count occurred at an average exponential decay rate of 0.149 day^-1^ for wild type (WT) and 0.179 day^-1^ for Δ*phaC* (see methods). After 20 days, WT cultures had retained around 20% of their original viability, whereas Δ*phaC* cultures had retained only around 4%. As seen in other organisms, the synthesis of PHB significantly affected long-term survival outcome.

**Figure 1.**
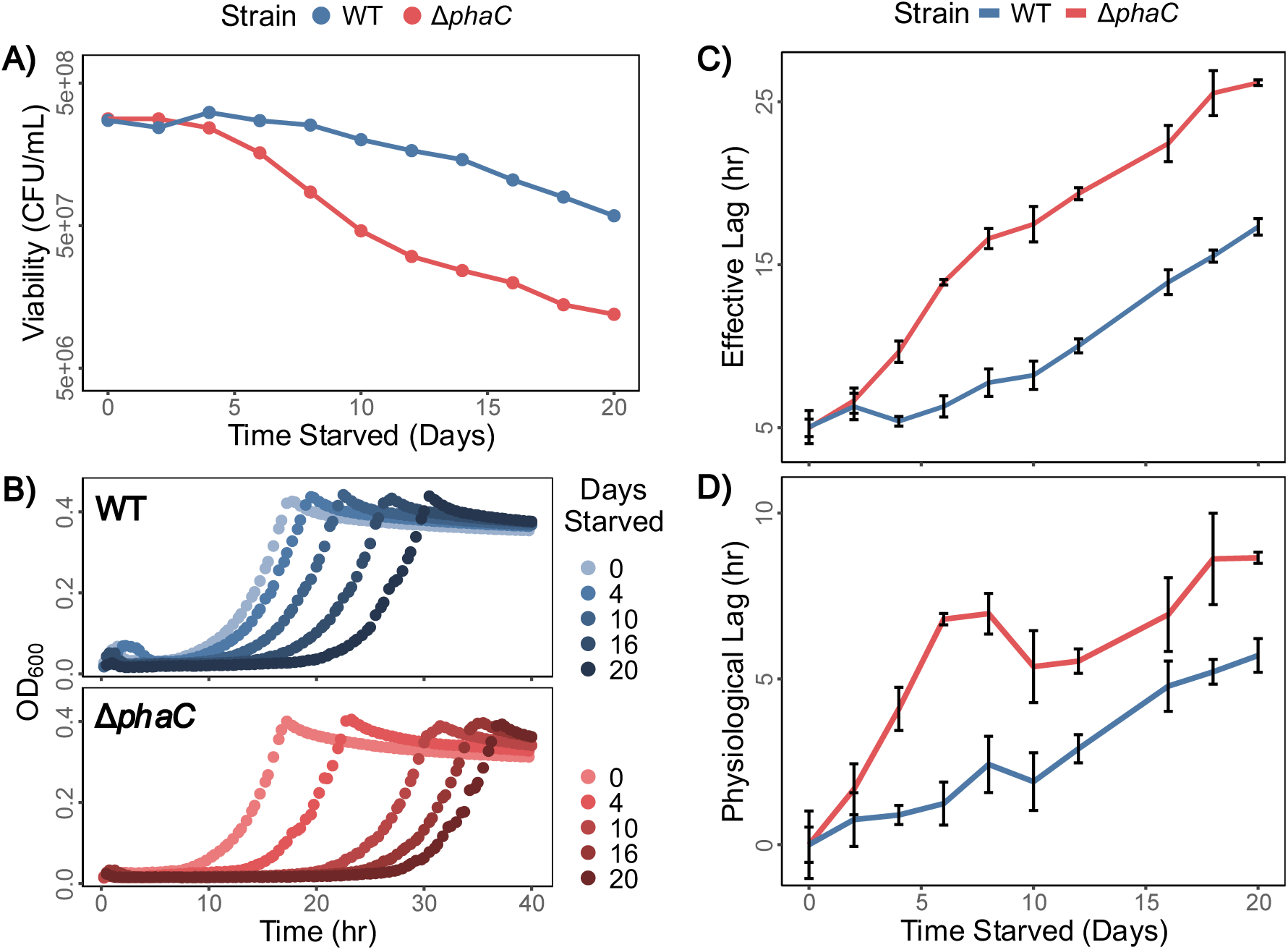
Loss in viability from starvation and accompanying population lag time in regrowth. For all parts, points represent the mean of three biological replicates; when present, error bars represent one standard deviation from the mean. (A) Culture viability measured by CFU/mL of WT and Δ*phaC* strains was assessed every 2 days by diluting and plating cells on agar. (B) Growth curves (OD_600_) of WT and Δ*phaC* strains that had starved for 0, 4, 10, 16, and 20 days. (C) Increase in effective lag as measured from growth curves. (D) “Physiological lag” of WT and Δ*phaC* strains as determined by the difference between calculated lag and simulated lag at a timepoint. Simulated lag was calculated from growth curves simulated from the mean number of CFUs present at that timepoint (See Fig. S1).

Complementary to tracking survival, we quantified lag (Fig. 1B) and growth rate (Fig. S2) upon reintroduction to media supplemented with succinate and ammonia. Any delay in regrowth observed by optical density (OD) occurs due to the joint effects of lowered viability (e.g. smaller starting population) and physiological lag of the surviving cells. Because of this, we denote the empirical measurement obtained from OD determined growth curves as the “effective lag” (see Fig. S1). While the effective lag for Δ*phaC* increased quickly within days of starvation, WT did not change significantly from non-starved for the first four days without nutrient supply (t-test, p = 0.229 for WT day 0 vs. 4 effective lag) (Fig. 1C). We also denote the difference between the effective lag of a starvation timepoint and the lag simulated from CFU at that time as the true “physiological lag” incurred due to starvation (see Fig. S1). We saw that having PHB metabolism influenced the “physiological lag” for surviving cells as well. The total physiological lag after 20 days was around 5 hours, or about 1.7 doublings for WT populations, while it was about 9 hours, or approximately 3 doublings for Δ*phaC* (Fig. 1D).

### Low osmolarity of starvation media positively impacts population outcome

Many other environmental factors also play significant roles in determining viability and lag. Since the osmotically balanced minimal medium we used does not reflect the full range of osmotic conditions encountered by natural microbes, we investigated the effects of variable osmotic pressure during starvation on viability and regrowth. We established three osmotic conditions during starvation: 1) low osmolarity, ultra-pure (18 MΩ-cm) water, 2) medium osmolarity, minimal media (1.88 mM Na^+^, 1.04 mM Cl^-^), and 3) high osmolarity, minimal media with an additional 100 mM NaCl added (see Methods). We starved populations in these osmotic conditions and assessed viability and regrowth every 5 days.

We discovered that out of the three conditions, cultures in high osmolarity lost viability the fastest regardless of PHB presence, with WT and Δ*phaC* undergoing around a 5.5- and 50-fold reduction in viability, respectively, after 20 days in starvation. (Fig. 2A). Surprisingly, populations which starved in the ultra-pure water maintained a viability insignificantly different from those in medium osmolarity (Linear mixed effects model, p > 0.1 for both WT and Δ*phaC*, low vs. medium osmolarity). Those populations also had a significantly shorter lag time than populations starved in medium osmolarity (Linear mixed effects model, p < 0.005 for both WT and Δ*phaC*, low vs. medium osmolarity) (Fig 2). Both strains showed comparable ratios of viability across the three conditions, suggesting the response to osmotic stress is independent of the use of PHB in starvation (Fig. 2A). Finally, we noted both a maintenance of viability and a plateau in lag time for all Δ*phaC* cultures between days 15 and 20 (Fig. 2).

**Figure 2.**
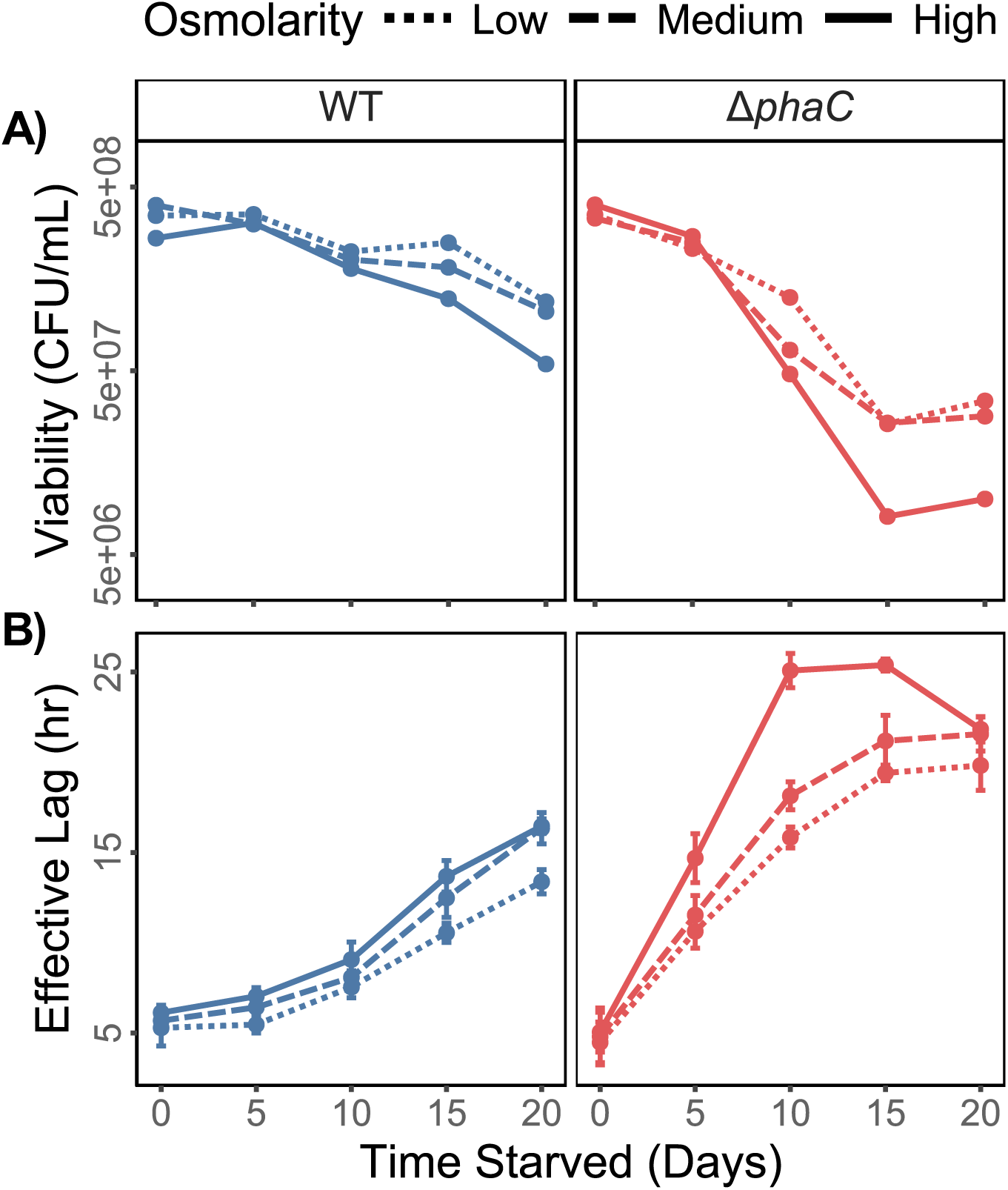
Loss in viability and an increase in effective lag for cells starving in a range of osmolarities. (A) Viability of WT and Δ*phaC* strains was assessed by diluting and plating cells on agar medium. Points represent the mean of three biological replicates. (B) Starved cells were reintroduced to fresh medium and effective lag quantified (see Fig. S1). Points represent the mean of three biological replicates and error bars one standard deviation.

### Heterogeneity in appearance time increases with time starved and reveals subpopulations

In addition to the population-level outcomes observed, we sought to quantify the variation in phenotypic traits such as lag time and growth rate within populations. When counting CFU to determine total viability, we observed that colony sizes were highly heterogeneous across all plates of cultures regrowing from starvation. We interpreted the distribution of colony sizes as a direct manifestation of heterogeneity in single cell lag time, as has been done in many previous studies on phenotypic variability (27, 43, 44). To quantify the observed heterogeneity, we developed an image analysis pipeline to track colonies growing on agar from their first optically detectable size (see methods). From this “trajectory” of colony size over time, appearance time and growth rate were extrapolated, giving us the distribution of critical phenotypic traits across the individuals in a population.

We starved WT and Δ*phaC* in media at the three osmolarities previously described and assessed how distributions of appearance time changed through starvation (Fig. 3A). We found that the difference in median appearance time between day 0 and 20 was minimal for all osmolarities, averaging around 6 and 9 hours for WT and Δ*phaC*, respectively. This was consistent with the physiological lag calculated from viability and effective lag (Fig. 1D).

**Figure 3.**
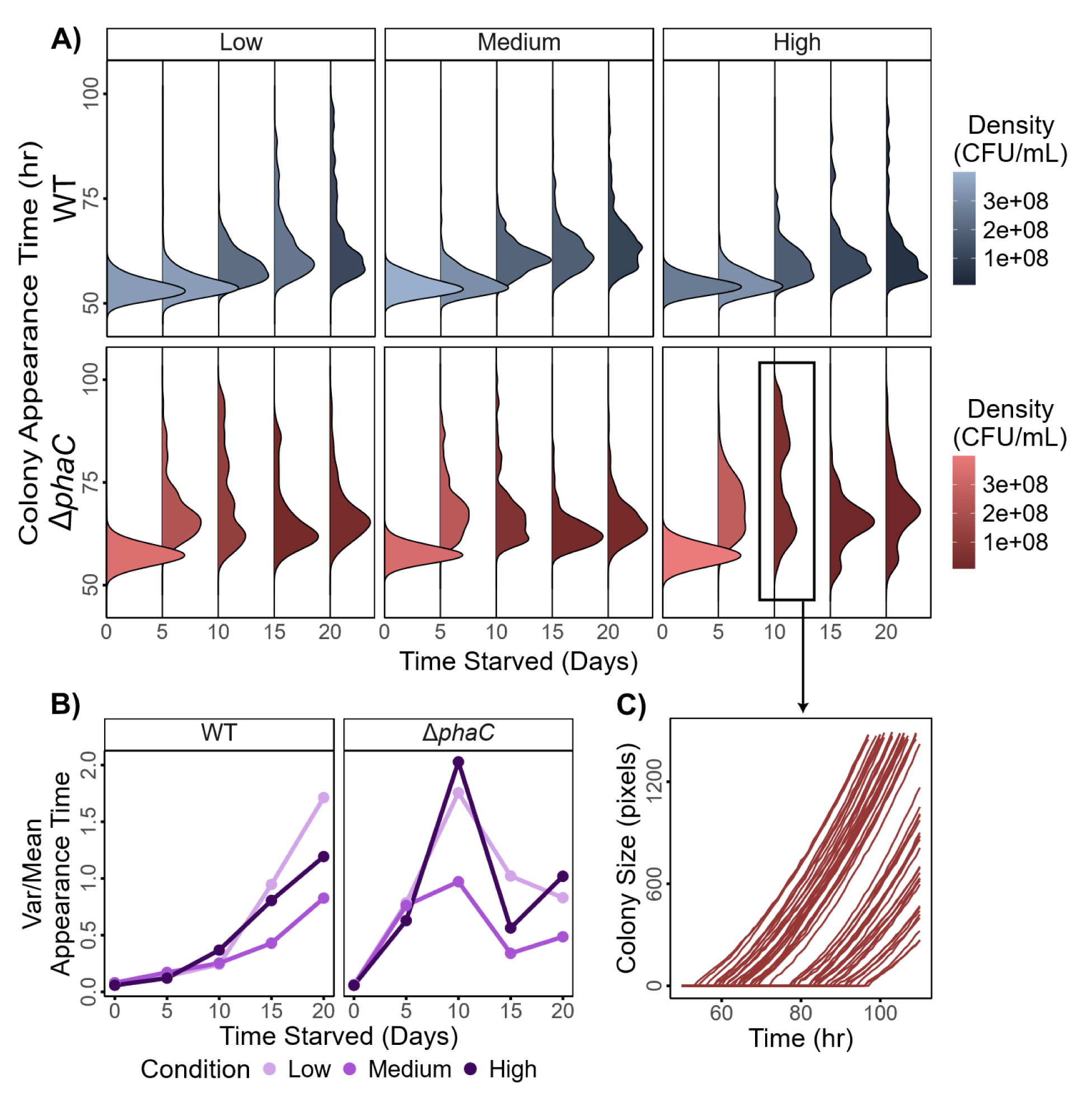
Colony appearance time distributions from 0-20 days in starvation for cultures in low, medium, and high osmolarity media. (A) WT (blue, top) and Δ*phaC* (red, bottom) colony appearance times in different osmotic conditions, colored by the density of the culture at that duration of starvation. Appearance was defined as the first time at which the colony regrowing on agar reached a size of 20 pixels when pictured at 800 dpi. Individual distributions represent the aggregate of data from three biological replicate cultures. (B) The variance-to-mean ratio of the colony appearance time distributions shown. (C) A random sample (n = 80) of Δ*phaC* colonies regrowing after starvation in high osmolarity for 10 days, displaying bimodality. Colony size trajectories (red lines) through time were obtained directly from image analysis.

In all strains and conditions, we observed a large skew towards late appearance times as starvation progressed. At high osmolarity, Δ*phaC* colonies showed a clear bimodal distribution of appearance time on day 10 with modes around 64 and 84 hours. The latter mode was absent at the next timestep (day 15), and the average time of the lower peak was then maintained for days 15 and 20, stabilizing around 10 hours later than the unstarved culture. The behavior of populations in all conditions appeared to be a manifestation of the same underlying process as for Δ*phaC* in high osmolarity, simply at a slower rate.

For all strains and conditions, variance in appearance time immediately increased from the non-starved distribution (Fig. 3B). In WT, this increase in variance occurred even before an increase in the median appearance time, and in general took the form of a skewed tail towards late appearance times. Additionally, the variance of colony appearance time did not correspond one-to-one with the population viability in either strain (Fig. 3B, 2A). The lowest variance in appearance time throughout starvation occurred at medium osmolarity, yet low osmolarity retained the most population viability.

### Osmotic effect on growth rate heterogeneity is absent in WT and highly variable in Δ*phaC*

In addition to appearance time, we extracted the distribution of growth rates from the colony trajectories. The formation of a colony from a single cell typically represents anywhere from 13 to 23 generations, depending on the organism, and colony specific growth rates have been found to be maintained throughout the entirety of the colonies growth (45). In liquid media, regrowth after starvation showed no significant deviation in growth rate from any one starvation timepoint to the next (Fig. S2). We therefore expected growth rate distributions to be consistent through time, potentially with increased variance. Instead, we saw a significant downward deviation in average WT growth rate between 0 and 5 days of starvation (t-test, p < 2.2 x 10^-16^ for day 0 vs. 5 across all conditions) (Fig. 4). This trend was consistent across all conditions yet not present in Δ*phaC*. To investigate whether the drop in average growth rate on day 5 was an artifact of the methods used to extrapolate growth rate, we compared WT colony trajectories at three timepoints (days 0, 5, and 10) for all osmotic conditions (Fig. S3). The slope of colony size vs. time was clearly different for 5 day colonies at any point in the trajectory, which showed that the lowered growth rate was not a product of the methods used.

**Figure 4.**
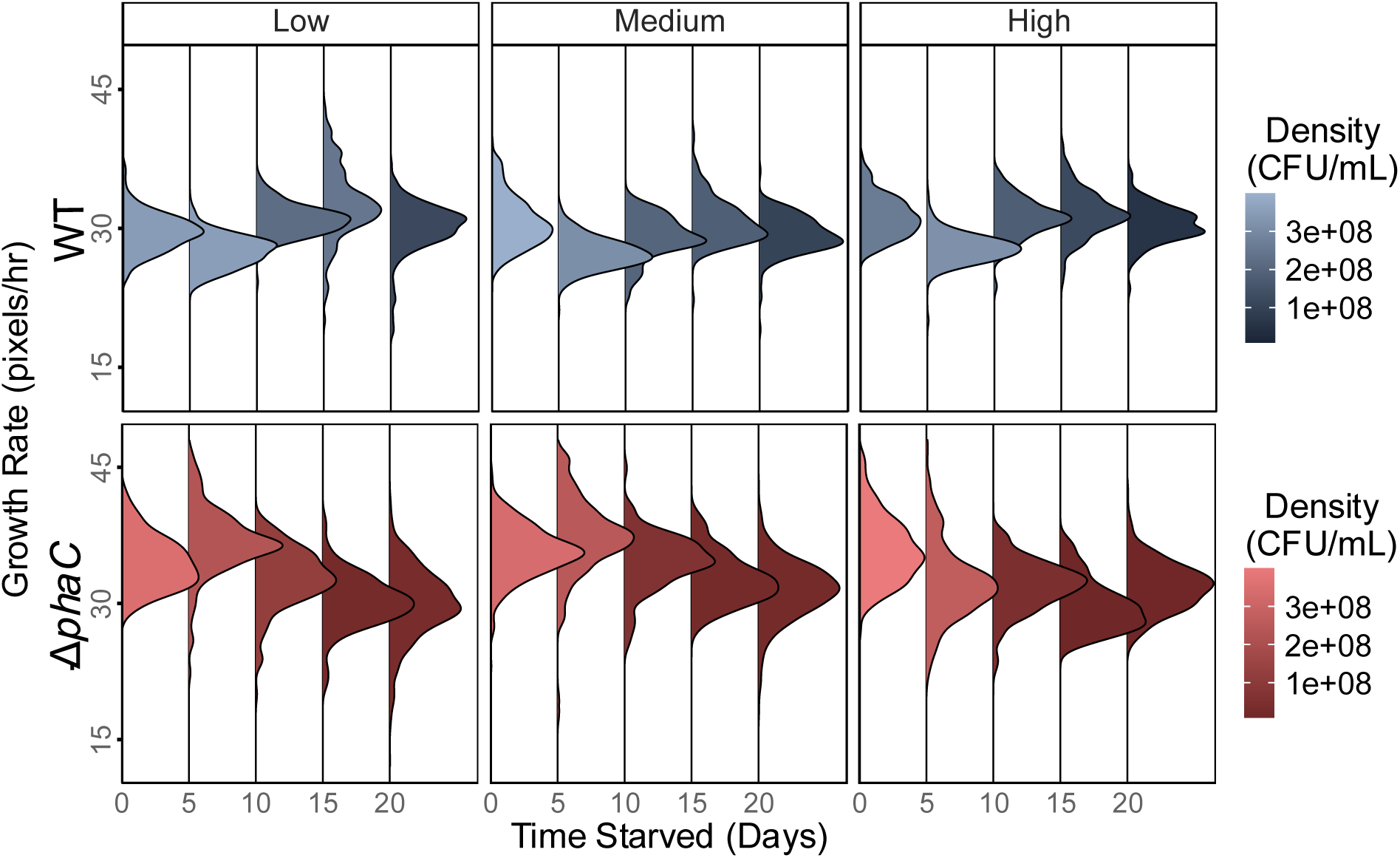
WT growth rates remain generally consistent while Δ*phaC* rates trend uniquely across osmolarities. WT (blue, top) and Δ*phaC* (red, bottom) growth rate distributions in a range of osmolarities, colored by the density of the culture at that duration of starvation. Individual distributions represent the aggregate of 3 biological replicates.

Δ*phaC* growth rate trends varied in multiple key aspects. For one, variance was consistently higher than in WT, even in unstarved cultures. Moreover, the median rate was significantly higher than WT in all conditions, until day 10 (Fig. S6). By day 20, the average growth rate between WT and Δ*phaC* had settled to roughly equal value. Importantly, the trends in Δ*phaC* median growth rate were unique across osmolarities and varied across a much larger range than for WT, with medians ranging from 37.4 to 29.7 pixels/hr (Fig. 4, S6).

## Discussion

In this work, we show that PHB metabolism positively impacts the survival of *M. extorquens* PA1 in starvation in three ways: 1) aiding cells in maintaining higher viability longer into starvation, 2) decreasing physiological lag in regrowth upon reintroduction to nutrients, and 3) decreasing variance in colony appearance time and growth rate. The beneficial effects of PHB have been documented in other closely related taxa, and the support in survival and regrowth seen here aligns with previous literature (15, 16). Remarkably, the magnitude of difference in loss of viability and lag seen between WT and Δ*phaC* here is likely due to a relatively small amount of accumulated PHB. WT cells grown at the 10:1 carbon-nitrogen ratio used in this experiment accumulate minimal PHB, nowhere near reported maxima of 60-80 percent of cell dry weight (12). Overall, these data reinforce PHB metabolism as a critical reservoir that stabilizes survival under energy deprivation, even under growth conditions where accumulation is minimal.

The benefits of PHB metabolism were not limited to one laboratory environment; across all osmotic conditions surveyed, death rate and lag times in Δ*phaC* cultures were consistently higher (Fig. 2). We found that a low osmolarity environment, namely ultra-pure water, allowed for greater maintenance of viability than in other osmolarities, regardless of PHB presence. This maintenance of over 25% of the original population after 20 days in starvation is markedly higher than many other bacteria for which starvation survival is reported (3). In related proteobacteria, *Bradyrhizobium* and *Cupriavidus*, ultra-pure water is the preferred medium for growth (21, 22). This proteobacteria specific abundance of hypoosmotic tolerance likely reflects the environments these species are isolated from, such as soil, leaves, oceans, and freshwater (46–48). However, the mechanisms allowing for survival and even growth in hypoosmotic media like ultra-pure water remain understudied (49). On the other side of the osmotic spectrum, additional salt in the starvation media provided a significant survival pressure, causing additional cell death as well as a wide variance in colony appearance time in regrowth (Fig. 2A, 3A). Tolerance to hyperosmotic stress has been identified in a broader region of the tree of life (20), and its molecular basis has been linked to the stringent response, membrane integrity, and oxidative stress response (50, 51). A preference for low osmolarity over high osmolarity might signify a lack of stress-related regulatory genes or a membrane more specialized to handle medium-to-low osmolarities.

The effects of osmolarity on viability in WT were directly proportional to that of the Δ*phaC* strain (Fig. 2A). This suggests that the underlying mechanisms for the response to osmotic stress and the benefit of PHB during starvation are distinct or uncoupled. Work with *Cupriavidus necator* showed that PHB granules provide a survival advantage to cells under hyperosmotic stress, but are not actively hydrolyzed to provide that advantage (52). Instead, granules initially became more liquid-like and “plugged holes” in the bacterial membrane to prevent cytoplasm leakage. Broader work has seen that general polyhydroxyalkanoate (PHA) production in mixed microbial communities increases in response to osmotic downshock (low osmolarity for short periods of time) (53). However, this study was done with actively growing cells. In our experiments, there was no opportunity for a growth phenotype to respond to osmotic stress, as the organisms were starving throughout the duration of exposure. Low osmolarity might still have induced a stress response in *M. extorquens*, but the response primarily increased variance in cell traits rather than decreasing the total viability (Fig. 3A, 3B).

The variance and bimodality we observed in colony appearance times after a duration of starvation (Fig. 3A) could represent a beneficial adaptation, or it could simply reflect the variance induced directly by stress. If the increased heterogeneity is an adaptation, it produced cells that assume both sides of the tradeoff between survival and growth: slower growing, resistant cells and faster growing, susceptible cells (Fig. S5). More work is necessary to determine the exact origin of colonies with distinct appearance times, but there are a few parallel examples that offer explanations. As seen in three γ-proteobacterial species (4), late-appearing colonies may represent cells that entered a dormant state due to stress from starvation or osmotic pressure, and take longer to regrow from dormancy than cells that remained somewhat metabolically active. Colony appearance distributions in Δ*phaC* in all conditions show tails towards late appearance times on days 5 and 10 in starvation, yet the tail is lessened on days 15 and 20 (Fig. 3A). A fraction of cells entering a dormant state might create the observed dynamic of certain colonies reappearing later and later after more time starved. Alternatively, as seen in *M. extorquens* with formaldehyde exposure (28), late-appearing colonies may represent cells that have accrued damage due to the stress of starvation. Those cells, on the verge of death before re-exposure to nutrients on agar plates then take longer to recover before resuming exponential growth, leading to long lag times.

The death of a less tolerant portion of the population would reveal a more tolerant portion that maintains viability and lower lag time. This would explain the colony appearance time distributions seen (Fig. 3). As cells less tolerant to starvation increased in lag time drastically (tails of distributions at late appearance time), cells more tolerant to starvation increase minimally in their lag time (5 to 9 hour increase over 20 days) (Fig. 3). This possibility also helps to explain the maintenance of viability and plateau in population lag time for all Δ*phaC* conditions between days 15 and 20 (Fig. 2), as later starvation timepoints would consist primarily of tolerant, fast-to-regrow cells. Recent studies focusing on stressful environments have discovered subpopulations tolerant to long-term nutrient depletion, labeled as GASP phenotypes (54, 55). These tolerant cells exhibit active metabolic profiles deep in starvation, scavenge for nutrients and other resources off of dead and senescent cells, and often accumulate mutations in global regulatory genes (54, 56–58). While proof of a GASP phenotype in Δ*phaC* would require further work, it is well supported by both the flattening of the death curve after day 15 (Fig. 2) and the maintenance of appearance times, most prominently in the early mode revealed in the bimodal distribution for Δ*phaC* in high osmolarity (Fig. 3).

We also observed an increase in the variance of appearance time for WT populations over the duration of starvation, but it was a monotonic increase as opposed to Δ*phaC*. Interestingly, within both WT and Δ*phaC*, low and high osmolarities had similar trends in variance despite having opposite survival outcomes (Fig. 3B, 2A). It is likely that both hypo- and hyper-osmotic environments present stresses to the populations, but only cells in low osmolarity were able to buffer the stress, leading to increased phenotypic heterogeneity. Non-lethal environmental shifts like nutrient-switches have been shown to induce variance, often preparing populations for future perturbations (24, 59).

Late-appearing colonies in longer durations of starvation also maintained lower growth rates. This trend was especially noticeable in populations regrowing from starvation in low osmolarity. This manifested as a negative correlation between growth rate and appearance time (Fig. S5). This follows logically from the idea that cells that grow more slowly form visible colonies later. However, the growth rate of the colony is measured from when it is first visible to many hours later, and colonies maintain that rate throughout. Depending on the microorganism, visible colonies typically represent 10^4^ to 10^7^ cells, or around 13 and 23 generations respectively since the cell that beget the colony was plated. This raises the question of why a maladaptive growth rate would be carried through roughly 20 generations. If it were genetic adaptation, the mutation would be required to be in the colony-forming cell, otherwise a deleterious change would be quickly outgrown by other colony mates that were not a mutant. Alternatively, it could be nongenetic memory of the stress of starvation passed on for many generations (60–63), thereby increasing the fitness of the population if starvation were to happen again.

Our results provide insight into how *Methylobacterium* persist in environments dominated by oligotrophy and osmotic fluctuation. Leaves, freshwater, and soil all impose cycles of feast, famine, and osmotic shock, where both consistent survival and the ability to exploit transient nutrient pulses are essential for population fitness (64). Cells with PHB metabolism are better poised to maintain viability, avoid slow-growing states, and regrow predictably when conditions improve. In contrast, the heterogeneity in appearance phenotypes seen in Δ*phaC* may represent maladaptive outcomes where timing and predictability are crucial for competition and colonization success.

## Materials & Methods

### Bacterial strains and culture conditions

The WT *Methylobacterium extorquens* PA1 (35) strain is CM2730, a cellulose deletion mutant (Δ*celABC*) that removed clumping. A Δ*phaC* strain lacking PHB, CM4916, was generated in the CM2730 background by two-step homologous recombination using pKB54 as a donor. This plasmid contains approximately 0.5 kb flanks upstream and downstream of *phaC* that were cloned into pPS04 (65) via a three-way Gibson ligation. The base medium used was a variant of *Methylobacterium* PIPES (MPIPES) minimal media (66) that lacked 5 mM (NH_4_)_2_SO_4_. This was also served as the “medium” osmotic condition. For the “low” osmolarity medium, we autoclaved 18Ω Ultra-Pure water. Alternatively, the “high” osmolarity was achieved by adding 100 mM NaCl to our base medium. Throughout, our medium for growth was base medium supplemented with 5 mM Na_2_SO_4_, 1.5 mM NH_4_Cl, and 3.75 mM sodium succinate (achieves a 10:1 carbon to nitrogen ratio). All liquid cultures were grown at a 5 mL volume in a 15 mL test tube with a loose lid, shaken at 250 RPM at a 45° angle, and incubated at 30 °C. For cultures on solid agar medium, standard MPIPES media (including the 5mM (NH_4_)_2_SO_4_) plus 16 g/L Bacto Agar was used, with 15 mM sodium succinate. Prior to all experiments, freezer stocks were streaked onto agar medium and allowed to grow for 4 days. Biological replicates were obtained via a single colony inoculating 5 mL liquid growth media for 48 hours.

### Starvation procedure

Liquid cultures at their final density (∼2×10^8^ CFU/mL) were transferred to 15 mL conical tubes and centrifuged at 4000 rpm for 10 minutes at 30 °C. The supernatant was discarded and cells were washed with 2 mL of starvation media. Tubes were vortexed to resuspend the cell pellet in the new liquid, and then centrifuged again at the same settings. Supernatant was again discarded and 5 mL of starvation media was added to the tubes. These tubes were then vortexed to resuspend the cell pellet and transferred to a new 15 mL test tube before incubation at 30 °C while shaking at 250 rpm.

### Viability analysis

All plating was performed from cultures that had already been transferred into starvation media as described in the section above. For each culture, 50 µL was transferred to 450 µL of starvation medium and mixed thoroughly. A 10-fold serial dilution series down to a total dilution factor of 10^-5^ was then performed and 50 µL of the chosen dilution was spread onto solid agar media. Most quantification was done at the 10^-5^ dilution, although some durations of starvation with fewer CFU required sampling at 10^-4^. Plates were then incubated at 30 °C, right side up if used for a colony tracking experiment. After 4-5 days of incubation, plates were stored at 4 °C until counting. To calculate the average rate of loss of viability from the data in Fig. 1A, the negative linear slope of natural logarithm transformed CFU/mL vs. time starved was calculated from the linear regression of all points after the last timepoint of non-significant change in CFU from unstarved culture.

### Growth analysis

To obtain growth curves, 10 µL of culture was inoculated into 630 µL of growth media in a (Corning) 48-well plate, incubated in a Biotek Synergy H1 Plate Reader (Agilent Technologies) at 30 °C, shaking at 548 rpm in a double orbital motion. Growth curves were obtained from raw plate reader data and analyzed with a custom R script (see Data Availability) employing the growthrates package. OD_600_ was blanked by subtracting from every data point the value of the minimum OD of the whole plate, and then adding 0.007 (∼1/64 of 0.5 OD, the OD of a typical inoculating culture). The growth rate was calculated as the maximum slope of the set of log-linear regression slopes that fit 6 hours of data. Effective lag was calculated as the time coordinate at which a horizontal line at the y-value of the first data point intersected with the log-linear regression line of maximum slope of the growth curve (see Fig. S1 for an example).

In order to distinguish the loss of viability from physiological lag in Fig. 1C, growth curves for cultures were simulated from the number of CFUs remaining at that timepoint that then followed the exponential growth equation, N(t) = N_0_*e^rt^, where N_0_ was the predicted inoculum in CFUs, r was the growth rate obtained from the corresponding culture’s empirical growth curve at that timepoint, and t was the time in hours post simulated inoculation. CFU values were then converted into OD_600_ using an OD_600_ vs. CFU standard curve generated in triplicate (see Data Availability). To obtain effective lag, OD_600_ values below 0.02 (the reported detection limit of the Biotek Plate Reader) were set as 0.02, and then lag was calculated using the growthrates package the same as for empirical data.

### Photobed scanner colony tracking

In order to follow variation in recovery between cells we used petri dishes into which 25 mL MPIPES Agar supplemented with 15 mM sodium succinate was dispensed. Cultures were diluted and spread across solidified agar at a concentration that yielded 100-300 expected colonies per plate. They were then incubated at 30 °C agar side down on Epson Perfection V600 Photobed scanners connected to a computer running Linux Ubuntu with the scanimage and temper utilities. Felt cloths were placed under plate lids to prevent condensation from interfering with photos and to provide a dark background. A custom batch script took 800 dot per inch (dpi) tiff photos of the plates and recorded the temperature at the time of the picture, scheduled hourly by a cron job. After 4-5 days of incubation, the time series of images was transferred to a python environment where colonies were labeled using a binary closing algorithm and a labeling function included in the scikit-image (67) image processing library. Colony pixel count was linked to metadata as a time series and output to a data frame that underwent filtering and analysis in R. From the time series of colony size, colony appearance time was defined as the first time at which the colony’s size was larger than 20 pixels, and growth rate was defined as the maximum slope of colony size in pixels vs. time in hours, for colony sizes between 50 and 500 pixels. To see the scripts used, see Data Availability.

## Data Availability

All raw data and code are deposited on GitHub.

https://github.com/keatjad/StarvationOsmolarity

## Acknowledgements

This work was supported by a National Science Foundation grant to CJM (DBI-2320667), a Department of Energy grant to CJM (DE-SC0022318), and by a Hill Undergraduate Research Fellowship from the University of Idaho to KJA. We thank Sean M. Caroll for generating the mutant strain Δ*phaC*.

