## Supplemental Figures for "Storage polymer polyhydroxybutyrate promotes recovery and limits phenotypic heterogeneity in regrowth from carbon starvation in *Methylobacterium extorquens*"

**Supplementary Information**

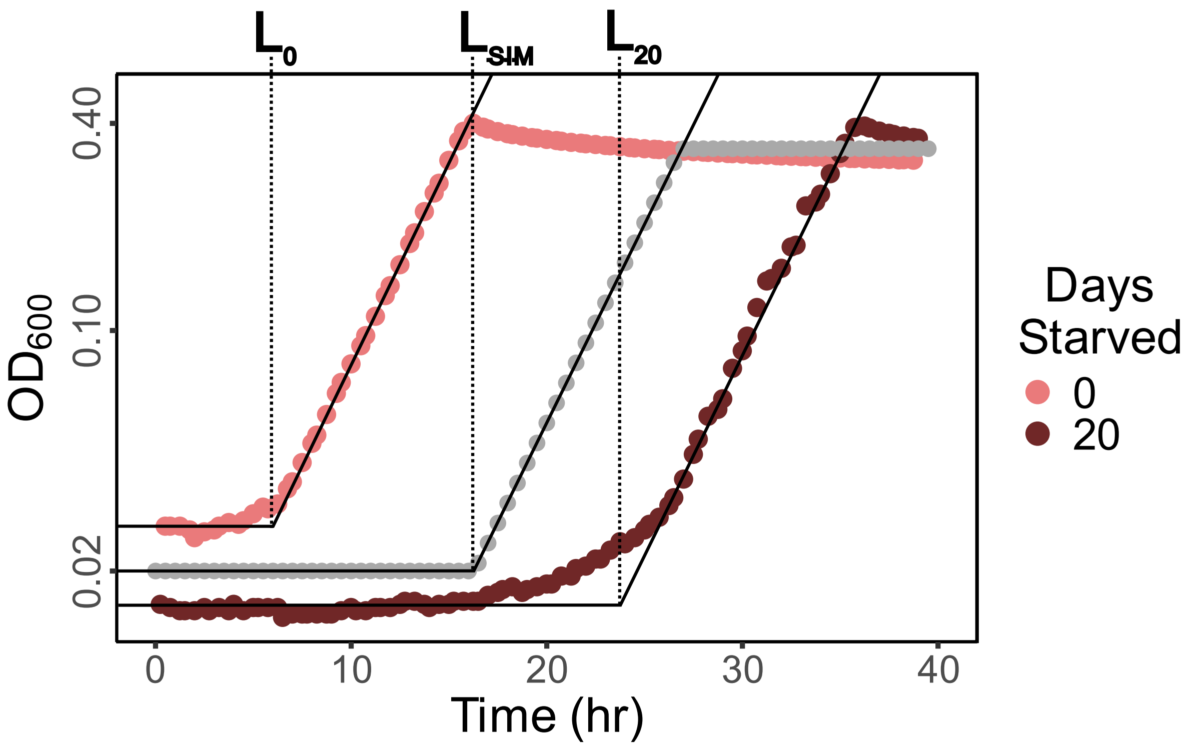


**Figure S1. Extraction of growth rate and effective lag from growth curves, where physiological lag is the difference between experimental and simulated growth curve.** The growth rate was the maximum slope of the linear fit of 6 hours of natural logarithm transformed $OD_{600}$. Effective lag was calculated as the time coordinate of intersection of the line defining growth rate and the horizontal line at a y-coordinate of the first $OD_{600}$ reading. Effective lag for Δ*phaC* cultures at days 0 and 20 of starvation are shown ($L_0$ and $L_{20}$). Simulated growth curve (gray) represents the expected $OD_{600}$ resulting from a culture inoculating growth media at the density of CFUs measured at day 20 of starvation. Physiological lag was calculated as the difference between effective lag and the simulated lag (in this example, $L_{20} – L_{SIM}$).

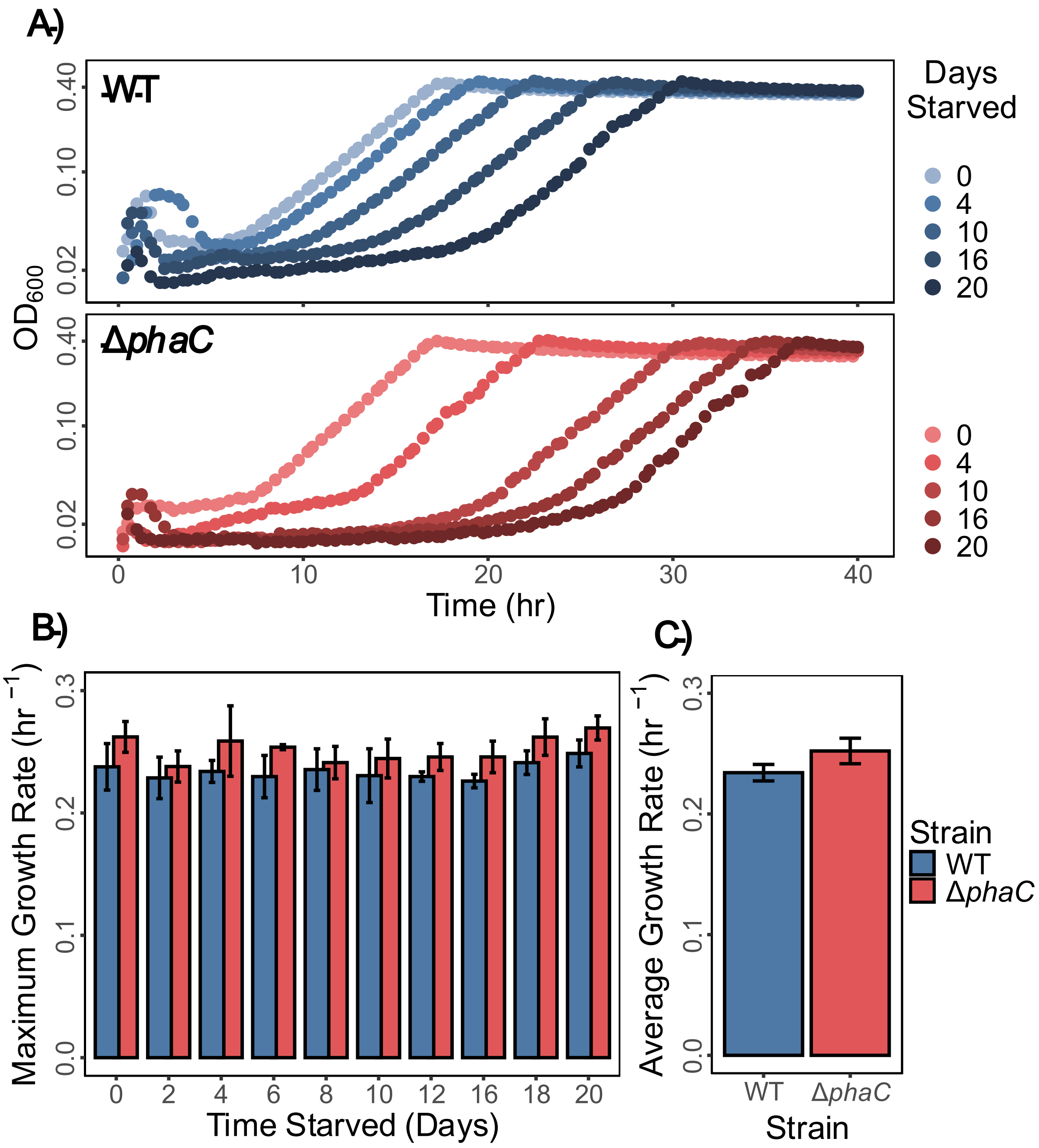


**Figure S2. Growth rates in regrowth from starvation show no significant deviation from non-starved populations in either strain.** (A) Natural log-transformed optical density ($OD_{600}$) of WT and Δ*phaC* regrowing from various durations of starvation show that while lag time varies, growth rate appears the same. (B) Growth rates extracted as linear fits of the data in part A show no significant deviation from directly adjacent timepoints across 20 days of starvation. Bars and error bars respectively represent the mean and 95% confidence interval of 3 biological replicates. (C) An average of growth rate from all time points in starvation show that Δ*phaC* has a higher overall growth rate. Bars and error bars respectively represent the mean and standard deviation of the 10 average growth rates obtained.

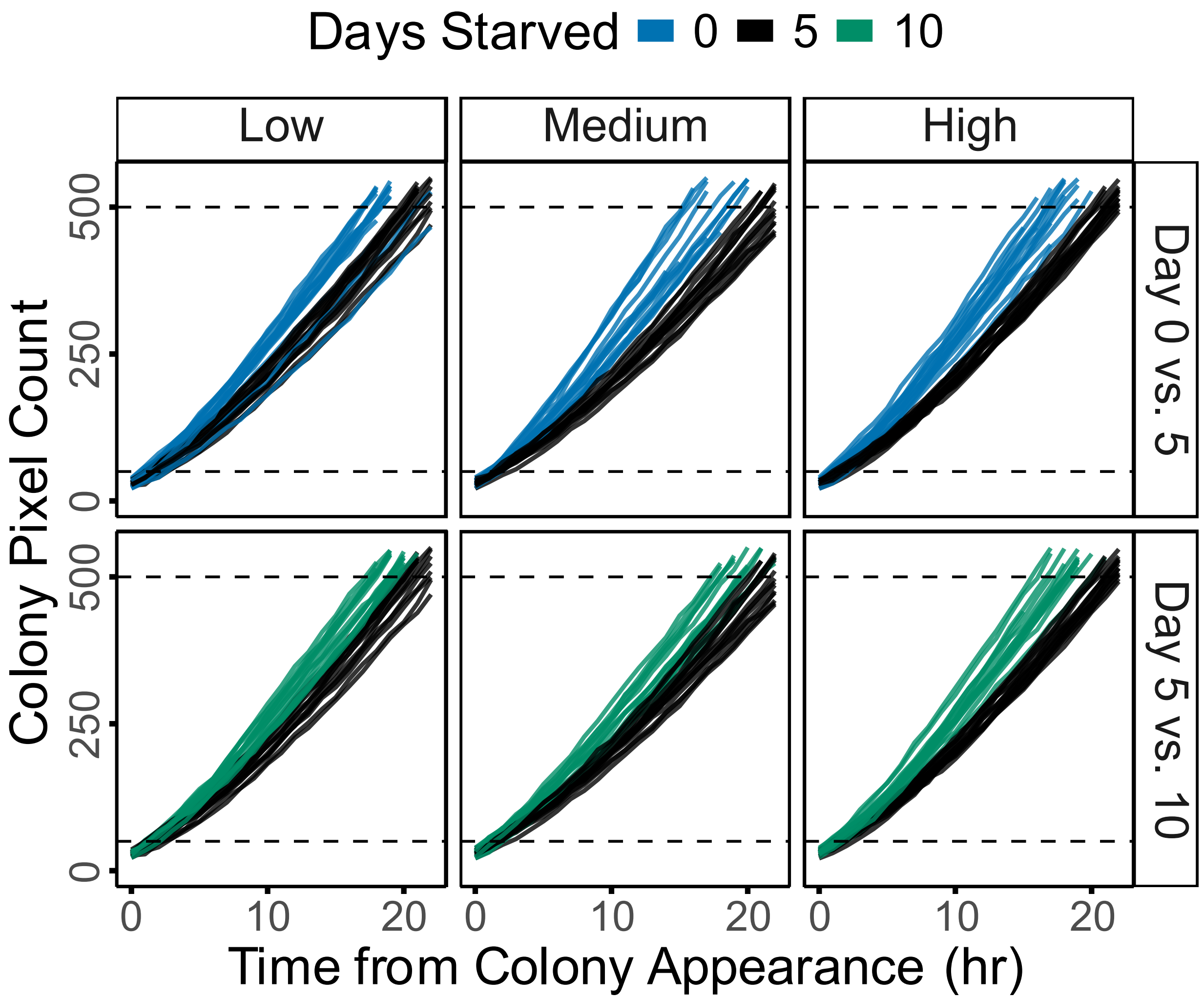


**Figure S3. WT 0 vs. 5 and 5 vs. 10 days starved colony trajectory comparisons display a significant decrease in growth rate for day 5 colonies in all conditions.** A random sample (N = 15) of colonies across three biological replicates represent colored traces shown, where the x-axis is the time since the colony became visible. Growth rate was calculated as the maximum slope of the linear fit of 5 colony pixel counts vs. time between the two horizontal dashed lines at 50 and 500 pixels.

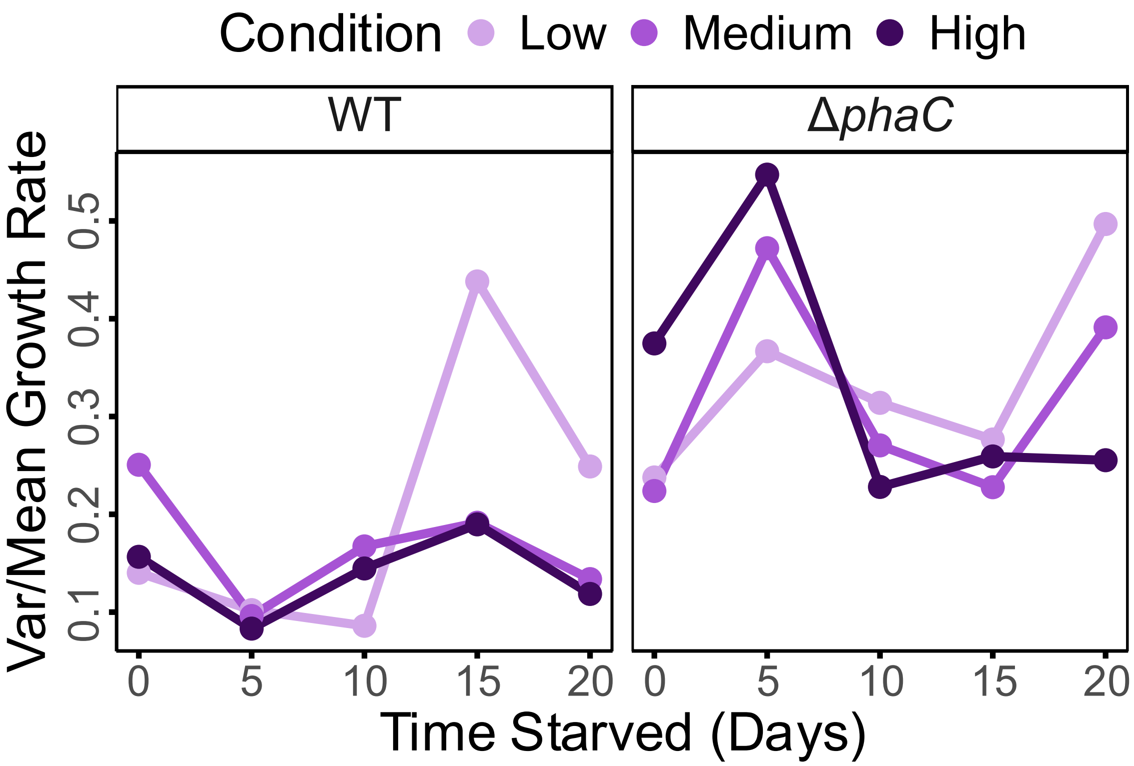


**Figure S4. The variance-to-mean ratio of colony growth rate distributions.** WT cultures (left) typically displayed a lower variance-to-mean ratio than Δ*phaC* cultures (right) in all osmotic conditions.

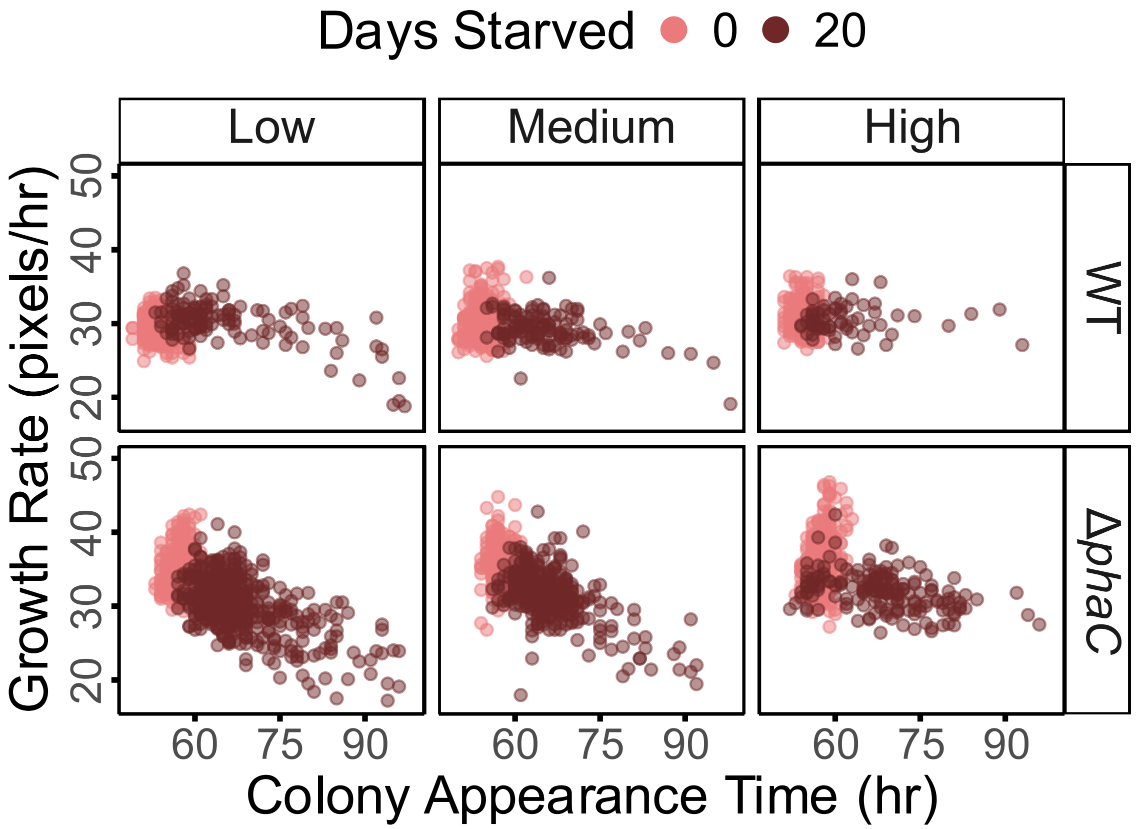


**Figure S5. Colony growth rates are negatively correlated with appearance times after 20 days of starvation in some conditions.** Correlation is weaker in all WT strains, as well as in high osmolarity for both strains.

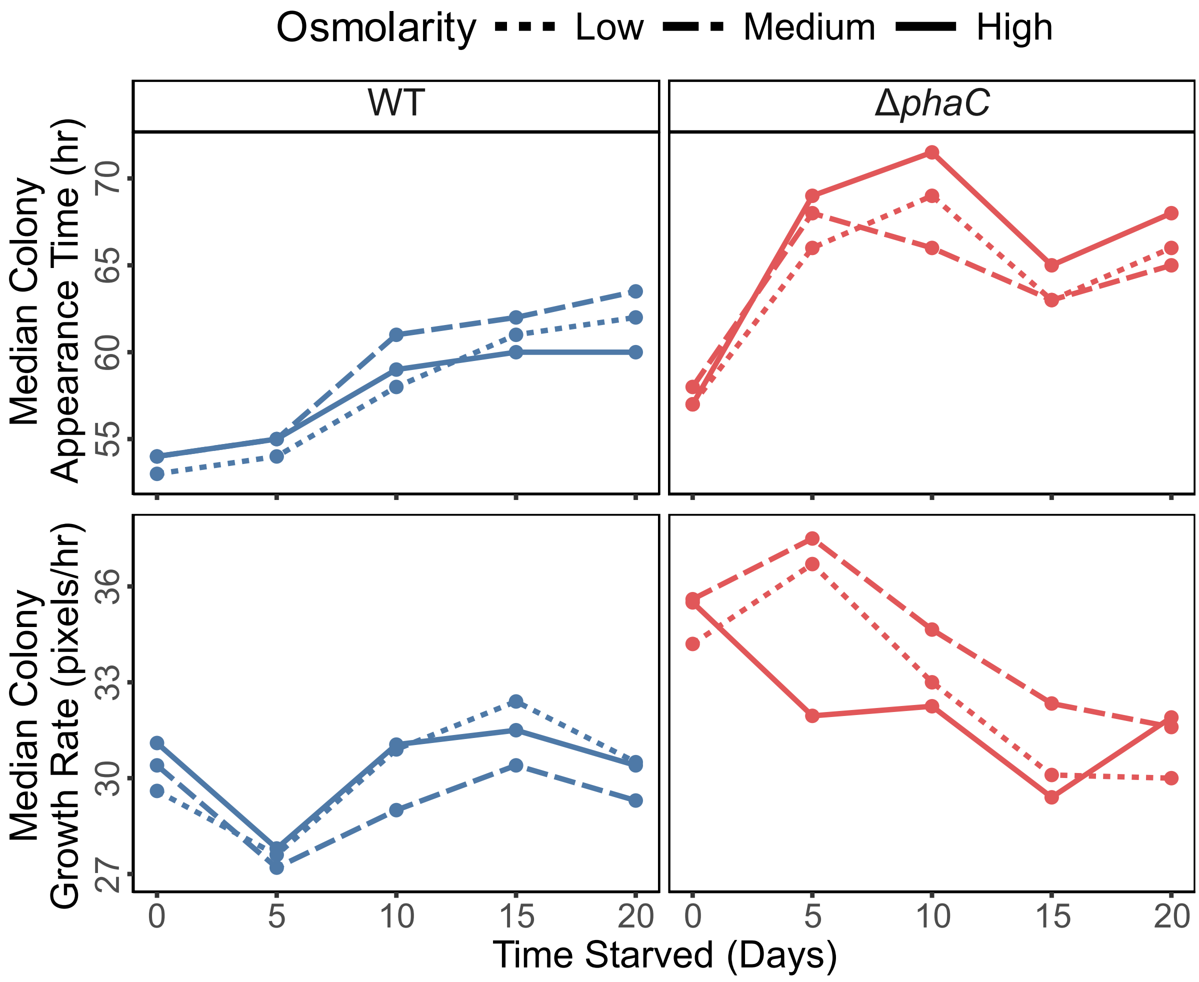


**Figure S6. Median of colony appearance time and growth rate distributions extrapolated from colony growth.** Median colony appearance time (top) increased monotonically in WT, but peaked and decreased in Δ*phaC*. Median colony growth rate (bottom) decreased on day 5 in WT, while it peaked on day 5 in Δ*phaC*. After 20 days of starvation, growth rates across strains approached the same value.
